# Simple shell measurements do not consistently predict habitat in turtles: a reply to Lichtig and Lucas (2017)

**DOI:** 10.1101/2024.03.25.586561

**Authors:** Serjoscha W. Evers, Christian Foth, Walter G. Joyce, Guilherme Hermanson

## Abstract

Inferring palaeoecology for fossils is a key interest of palaeobiology. For groups with extant representatives, correlations of aspects of body shape with ecology can provide important insights to understanding extinct members of lineages. The origin and ancestral ecology of turtles is debated and various shell or limb proportions have been reported to correlate with habitat ecology among extant turtles, such that they may be informative for inferring the ecology of fossil turtles, including early shelled stem turtles. One recently described method proposes that simple shell measurements that effectively quantify shell doming and plastron width can differentiate habitat classes among extant turtles in linear discriminant analysis, whereby aquatic turtles have low domed shells with narrow plastra. The respective study proposes unorthodox habitat predictions for key fossil turtles, including aquatic lifestyles for the early turtle *Proganochelys quenstedtii* and the meiolaniform *Meiolania platyceps*, and terrestrial habits for the early turtle *Proterochersis robusta*. Here, we show that these published results are the consequence of questionable methodological choices such as omission of species data which do not conform to a preconceived shell shape-ecology association. When these choices are reversed, species measurements for fossils are corrected, and phylogenetic flexible discriminant analysis applied, habitat cannot be correctly predicted for extant turtles based on these simple shell measurements. This invalidates the method as well as the proposed palaeohabitats for fossils.

## Introduction

Inferring the palaeoecology of fossil species is of central importance for the field of palaeobiology, as knowing the ecological attributes of organisms (e.g., habitat or diet) allows researchers to test if or how evolutionary patterns in the origin of lineages and body plans are related to ecology. For turtles, habitat ecology has been discussed to be important as drivers of their biogeographic distribution (e.g., Joyce et al. 2016; Ferreira et al. 2018), body size evolution (Farina et al. 2023), ecomorphological diversification (e.g., Hermanson et al. 2022), body shape and proportions (e.g., Hermanson & Evers 2024), dietary adaptations (e.g., Claude et al. 2004; Foth et al. 2017; Ponstein et al. 2024), morphological and functional innovations related to locomotion (e.g., Joyce & Gauthier 2004; Evers et al. 2019), and also the origin of the shell as the most characteristic trait of turtles (e.g., Rieppel & Reisz 1999; Rieppel 2013; Lyson et al. 2016; Schoch et al. 2018). However, the habitat ecology of fossil turtles can be difficult to know, for example when allochthonous fossil deposition may be invoked for turtles found in aquatic depositional environments (e.g., *Odontochelys semitestacea*: Li et al. 2008; Joyce 2015; thalassochelydians in Solnhofen lagerstätten deposits: Anquetin et al. 2017; Joyce et al. 2021). Researchers frequently try to synthesize simple anatomical observations that reliably (i.e., accurately and precisely) correspond to (habitat) ecology among extant turtles, proposing that these can be used to ecologically classify extinct turtles (e.g., Joyce & Gauthier 2004; Dudgeon et al. 2021). Lichtig and Lucas (2017) recently proposed a method that allows inferring the habitat palaeoecology (i.e., aquatic versus terrestrial) of fossil turtles based on simple shell measurements. The underlying observation is one that has long been observed: aquatic turtles on average have flatter shells than terrestrial turtles, whereby flatness is commonly interpreted as a hydrodynamic adaptation whilst a high domed shell morphology can aid in self-righting (e.g., Romer 1967; Claude et al. 2003; Domokos & Várkonyi 2008; Rivera 2008; Benson et al. 2011; Stayton 2011; Polly et al. 2016; Williams & Stayton 2019; Stayton 2019; Ferreira et al. 2024). Although there are of course gradients of “aquaticness” among turtles (e.g., with many testudinids never entering a body of water, many chelonioids and trionychids only leaving the water to lay they eggs, but some turtles, such as the wood turtle *Glyptemys insculpta* readily spending time in water or on land [Ernst and Barbour 1989]) that could be further anatomized, the principal distinction between terrestrial and aquatic species is a meaningful categorization, for several reasons. First, one of the most important functional aspects of aquatic lifestyles is a habitual submersion in water (e.g., Fabbri et al. 2022b), which can occur for different reasons, including foraging or seeking protection. Aquatic animals face functional challenges that are different from functional challenges imposed on animals that never submerge (e.g., Joyce & Gauthier 2004, Fabbri et al. 2022b). Evidence for this among turtles comes, for instance, from differences in hand structure in aquatic bottom walking chelydroids and terrestrially walking tortoises (Joyce & Gauthier 2004). For the purpose of shell geometry, there is a clear expectation formulated in Lichtig and Lucas (2017), but also in other studies (e.g., Stayton et al. 2018), that shell geometry is influenced by hydrodynamic adaptations, which should universally apply to turtles that enter the water and universally be absent to those that do not. Secondly, the clear definition and distinction of variables is beneficial to analyses of ecomorphology (e.g., Fabbri et al. 2022b), and the binary distinction of turtles that never enter the water (i.e., terrestrial turtles) and turtles that do (i.e., aquatic turtles) provides a clearly testable habitat hypothesis that is not confounded by the varying degrees of aquaticness. Lastly, the principal categorization into aquatic and terrestrial turtles is one that has been used frequently in ecomorphological studies (e.g., Claude et al. 2003; Benson et al. 2011; Stayton 2011; Wise & Stayton 2017; Stayton et al. 2018), and even in those that also use finer degrees of classifying aquatic taxa (e.g., Joyce & Gauthier 2004; Foth et al. 2017; Hermanson et al. 2022; Evers et al. 2022). As such, this categorization is useful for literature comparisons of other studies focused on shell geometry but also those that study other anatomical systems influenced by aquatic/terrestrial adaptations, such as limbs (e.g., Joyce & Gauthier 2004). Lichtig and Lucas (2017) used biplots and discriminant analysis of measurements obtained from extant turtles, in particular sagittal doming (i.e., carapace length to shell height ratio) versus plastral width (i.e., carapace width to plastron width ratio), to establish that shell ratios can discriminate between aquatic and terrestrial turtles. Cursory examination of the figured plot (their figure 3), however, reveals some oddities. For instance, the fossil stem-trionychian *Basilemys gaffneyi* plots as one of the highest-domed turtles, although this species is known for its flat shell morphology (Sullivan et al. 2013). This casts doubt on the primary data of Lichtig and Lucas (2017), which led us to examine their data in more detail to validate their study.

We, here, show that the published dataset of Lichtig and Lucas (2017) has errors, that their statistical analysis exhibits fundamental flaws, and that a corrected phylogenetic statistical analysis of their data cannot support the conclusion of these authors, namely that simple shell measurements can accurately and reliably predict the ecology of living and fossil turtles.

## Material and Methods

### Illogical fossil measurements in the original data and newly measured data

The tables and supplementary data of Lichtig and Lucas (2017) contain illogical values for a number of turtles. For instance, the plastron widths and carapace lengths of all taxa given in their table 1 exceed the values for carapace width and carapace length, respectively – which is anatomically impossible for the former and not realized among any known extant or fossil turtle for the latter. This is likely a result of data having been entered into the wrong column, but the columns do not seem to have simply been mislabeled, because simply rearranging the data columns also do not lead to plausible results. For example, even if the values provided for *Basilemys gaffneyi* (USNM 11084) are arranged into a sequence that makes sense for the morphology of the specimen (i.e., carapace length > carapace width > plastron width > shell height; see Sullivan & Lucas 2015), the datapoint falls at an average doming value for turtles, which contradicts expectations for the species. To produce more accurate values for the genus *Basilemys*, we took novel measurements of four specimens, following the measurement instructions of the original paper (i.e., Lichtig & Lucas 2017). We computed new shell ratio measurements based the 3D model of a specimen of the species *Basilemys morrinensis* (https://www.morphosource.org/concern/media/000031637) that was described by Mallon and Brinkman (2018). This specimen (CMN 57059) is likely somewhat compressed dorsoventrally, but the original description states explicitly that the low shell profile of this specimen is consistent with other, relatively uncrushed *Basilemys* shells (Mallon & Brinkman 2018), which is an observation that we agree with. We also added measurements for a specimen of *Basilemys variolosa* (CMN 8516) based on figures by Langston (1956). This specimen is also likely somewhat compressed, but has also been reported to be relatively undistorted (Mallon & Brinkman 2018). As a third specimen, we added AMNH FARB 5448, a specimen listed as *Basilemys* sp. by the AMNH but which has been referred to *Basilemys variolosa* (e.g., Brinkman 2003). The specimen includes a nearly perfectly preserved 3D shell with no apparent taphonomic flattening. The shell is available as a 3D file produced with surface scanning by the AMNH, and provided on the online repository MorphoSource, from which we downloaded it (https://www.morphosource.org/concern/media/000611825). As a fourth specimen of *Basilemys*, we downloaded a 3D model of DMNS EPV 103391-4891, *Basilemys* sp., which is also available on MorphoSource (https://www.morphosource.org/concern/media/000623817).

**Table 1.** Measurements of key ratios as provided in the supplementary data of Lichtig and Lucas (2017: LL17) and our corrected values. Note that the values for LL17 do not match their table 1 in all instances, as their supplements and tables do not agree. Also note that we use a specimen of *Basilemys morrinensis* and two specimens of *Basilemys variolosa* instead of *Basilemys gaffneyi* (see methods).

| Species (specimen used for ratio; source study) | Carapace length–height ratio (sagittal doming) | Carapace width–plastron width ratio | Carapace width–height ratio (coronal doming) |
| --- | --- | --- | --- |
| *Proganochelys quenstedtii* (SMNS 10012 and 17204; LL17) | 2.1 | 2.1 | NA |
| *Proganochelys quenstedtii* (SMNS 16980; this study) | 2.6 | 1.84 | 2.79 |
| *Proterochersis robusta* (SMNS 17561; LL17) | 1.39 | 1.94 | NA |
| *Proterochersis robusta* (SMNS 17561; this study) | 2.0 | 2.1 | 1.86 |
| *Meiolania platyceps* (AM F:57984 and 18775; LL17) | 2.0 | 2.54 | NA |
| *Meiolania platyceps* (AM F:57984 and 18775; this study) | 2.0 | 2.3 | 1.55 |
| *Basilemys gaffneyi* (USNM 11084; LL17) | 1.57 | 1.45 | NA |
| *Basilemys morrinensis* (CMN 57059; this study) | 4.26 | 1.64 | 3.32 |
| *Basilemys variolosa* (CMN 8516; this study) | 4.34 | 1.55 | 4.2 |
| *Basilemys variolosa* (AMNH FARB 5448; this study) | 3.86 | 1.62 | 2.82 |
| *Basilemys* sp. (DMNS EPV 103391-4891) | 4.33 | 1.63 | 3.53 |

Other fossil measurement data of Lichtig and Lucas (2017) exhibit similar, non-replicable ratio values, including those for *Proterochersis robusta* (SMNS 17561) and *Proganochelys quenstedtii* (based on multiple specimens). Indeed, some of the values provided in table 1 of Lichtig and Lucas (2017) mismatch those from the supplements (e.g., *Proterochersis robusta* carapace length-height ratio is listed as 1.82 in their table 1 but as 1.39 in the supplements and figure 3). Neither the ratios from table 1 nor the ones in the supplements could be reproduced by us when using the same data sources as listed in Lichtig and Lucas (2017). Thus, we repeated the measurements of Lichtig and Lucas (2017) for *Proterochersis robusta* on the same data source (images in Szczygielski & Sulej 2016) as well as on a 3D model of the specimen (SMNS 17561; Dziomber et al. 2020; available at: https://www.morphosource.org/concern/media/000116495) to produce measurements that reflect the dimensions of the actual specimen (see Table 1). These two versions of measurements were broadly in agreement, but as we deem the measurements on the 3D model more precise and easier to reproduce, we used our measurements based on the 3D specimen for all the analyses below. We also retrieved alternative measurements for *Proganochelys quenstedtii* (Table 1) based on a 3D model of SMNS 16980 (Dziomber et al. 2020; available at: https://www.morphosource.org/concern/media/000116494). We could, however, verify the shell ratios of Lichtig and Lucas (2017) for *Meiolania platyceps* based on figures in Gaffney (1996; a reconstruction based on AM:F 57984 and 18775; see criticism in Brown & Moll 2019), and differences between our and their values for this species are within the range of expected measurement error when basing ratios on published figures.

We added our novel measurements to the supplementary data of Lichtig and Lucas (2017), but we did not systematically cross-check the measurements for extant species, as these appear plausible. However, we added measurements for four chelonioid specimens belonging to the two sea turtle species that were previously included in the dataset of Lichtig and Lucas (2017; i.e., *Eretmochelys imbricata*, *Chelonia mydas*). The original measurements for these species had no recorded data for the height of the shell, which we used to calculate a coronal doming measure (see below). Thus, we added two specimens of *Chelonia mydas* (YPM VZ018243; SMF 73551) and *Eretmochelys imbricata* (SMF 36067; one unnumbered SMF specimen) to our dataset.

For plots and analyses, we deleted other fossil turtles with implausible values from the dataset (i.e., *Adocus bossi*, *Denazinemys nodosa*, *Scabremys ornata*, *Thescelus hemispherica*), as we were both unable to re-arrange the values into a meaningful order, but also unable to obtain new measurements. Thus, our dataset includes only five fossil species, for which one species (*Basilemys variolosa*) is represented by two specimens. To graphically show how measurement corrections for fossils result in different placements in the shell doming vs. relative plastral width biplots, our datasheet retains the original values for five fossils from the supplements of Lichtig and Lucas (2017). In addition, the dataset contains the original measurements taken from 213 individuals of 94 extant species of Lichtig and Lucas (2017) (taxonomy following the TTWG 2021) plus the new measurements of fourc chelonioids. Thus, our final, corrected dataset includes 217 individuals of 94 extant species (taxonomy following the TTWG 2021) alongside the above-mentioned fossil data (eleven datapoints, of which four are original measurements by Lichtig & Lucas 2017, and seven are new/corrected measurement data).

### Unjustified exclusion of datapoints and correction of habitat ecologies

Lichtig and Lucas (2017) mention that they discarded several extant terrestrial taxa with relatively flat shells from their plots and analyses because they interpreted them as “outliers” (Lichtig & Lucas 2017: p. 4). However, the affected species (i.e., *Malacochersus tornieri*, the pancake tortoise, and *Homopus boulengeri*, Boulenger’s cape tortoise) are valid biological observations of shell shape and we thus argue that their exclusion was unjustified. Lichtig and Lucas (2017) suggested that increased shell flexibility in these turtles justifies their exclusion, as shell flexibility causes varying shell heights. However, the “inflation” tactics used by *Malacochersus tornieri* to lock themselves into crevasses is achieved by soft tissue inflation through an enlarged central plastral fontanelle (Moll & Klemens 1996) and interpulmonary pressure is not sufficient for true shell inflation (Ireland & Gans 1972). The osteological flexibility of the shell indeed seems minimal in the absence of true joints between carapacial bones (e.g., NHMUK 1969.2211-2212; UF H85285). Thus, although the overall shape of *Malacochersus tornieri* is certainly unusual for a terrestrial tortoise, we see no well-justified osteological reasons to omit this species a priori, especially if the scientific question is specifically related to osteological measurements and their relation to ecology. For *Homopus boulengeri*, we are not aware of any literature that proposes that this species has a flexible carapace, even if species of the genus *Homopus* are generally comparatively flat-shelled tortoises. Consequentially, we re-inserted both species into our primary dataset.

We also assigned habitat ecologies to *Rhinoclemmys areolata* (i.e., terrestrial) and *Rhinoclemmys funerea* (i.e., aquatic) (Ernst & Barbour 1989), which were previously scored as “not assigned” for their ecologies in the supplements of Lichtig and Lucas (2017). We furthermore corrected the habitat preference of *Malacochersus tornieri* in the data spreadsheet from “aquatic” to “terrestrial”. Using this corrected dataset, we reproduced versions of the original plot of Lichtig and Lucas (2017: their figure 3) based on specimen-level data as well as species means.

Comparison of our sagittal doming plots with the figure 3 of Lichtig and Lucas (2017) suggests that at least one aquatic turtle species is absent from their plot although it is part of the supplementary file that forms their data basis. As only about 100 data points can be discerned from their plot, we presume it shows species means for taxa with multiple measurements, whereas their figure caption suggests that the data correspond to specimens, of which there should be 225 according to their supplements (250 according to their figure caption). Nonetheless, one aquatic turtle seems to have been omitted from the plot completely. Specifically, the mean species values for the aquatic box turtle (*Terrapene coahuila*) are 2.1 for the carapace length to height ratio and 1.29 for the carapace width to plastron width ratio, but no aquatic datapoint is found at these coordinates. Indeed, as this data point should be located within the area of the graph only occupied by terrestrial species, we suspect that it was omitted as yet another “outlier”. This is not reported as such, although the values for *Terrapene coahuila* are discussed as “odd” and “inaccurate” (Lichtig & Lucas 2017: p. 6). However, the values recorded in the supplements of Lichtig and Lucas (2017) for *Terrapene coahuila* are broadly consistent with the species’ morphology and shell measurements provided by other studies (e.g., Burroughs et al. 2013), such that we included them into both our plots and comparative analyses (below).

We additionally created a plot that uses coronal doming (i.e., carapace width divided by carapace height) as an alternative doming variable, as this is the more typical measure of doming used in the turtle literature (e.g., Benson et al. 2011; Dziomber et al. 2020).

### Re-analysis of corrected original data

Lichtig and Lucas (2017) performed a linear discriminant analysis to test if extant species can be correctly classified as “aquatic” or “terrestrial” based on their shell measurements. However, as noted above, they removed numerous “outliers” from their analysis. Besides the fact that these species should not have been removed from analysis, we note that the appropriate statistical treatment of species data are phylogenetic comparative methods due to phylogenetic autocorrelation (Felsenstein, 1985). Here, we performed phylogenetic flexible discriminant analysis (pFDA; Motani & Schmitz 2011) on the relationship of sagittal as well as coronal doming and relative plastral width using our measurement-corrected version of the Lichtig and Lucas (2017) dataset. We performed this analysis twice, once on the full dataset and once with a reduced dataset that excludes the three “outlier” species mentioned above, *Malacochersus tornieri*, *Homopus boulengeri* and *Terrapene coahuila*. Although we do not follow the arguments for the exclusion of outliers, we performed the second analysis as we acknowledge that opinions on this may differ, and as this can show that discrimination between aquatic and terrestrial turtles is not possible based on the proposed simple shell measurements even when the strongest “outliers” (i.e., turtles that do not conform to an “aquatic species have low domed shells” association) are disregarded. All analyses were done in the R statistical environment (R Core Team 2021).

pFDA allows the test of whether a predictor (here, the relationship of shell measurement ratios) can correctly discriminate between known ecologies of a training dataset. Thus, the training dataset only includes extant species. If the training dataset is successful in discriminating between ecologies, pFDA can further be used to predict unknown ecologies based on the predictor (Motani & Schmitz 2011, Close & Rayfield 2012; Angielczyk & Schmitz 2014; Choiniere et al. 2021; Fabbri et al., 2022a; Hand et al. 2023; Leavey et al. 2023; Lowi-Merri et al. 2023; see Foth et al. 2019, Stayton 2019, and Hermanson et al. 2022 for turtle examples), such as palaeoecology from fossil shell measurements. The analysis thus basically tests if simple shell ratio measurements can indeed be used to predict habitat ecology of turtle species.

Phylogenetic comparative methods require a time-scaled phylogeny. We used trees from Thomson *et al*. (2021), who provided a posterior sample of 100 trees from their Bayesian node-dating analysis of molecular sequence data, which represent different time-scaled trees of the same topology. We used the “bind.tip” function of the ape v.5.0 package (Paradis & Schliep 2019) and the “date_nodes” function of Claddis 0.6.3 (Lloyd 2016) to add the five fossil species using a minimum branch length argument of 1 Ma (Laurin 2004) and published stratigraphic ages of the fossils. Hereby, *Proterochersis robusta* is used as the earliest branching testudinatan from our sample, following phylogenetic studies that include the latest anatomical insights from *Proterochersis robusta* (Szczygielski & Sulej, 2016, 2019). *Proganochelys quenstedtii* and *Meiolania platyceps* were added as successively more crownwardly placed stem turtles, following the consensus topology across multiple studies (e.g., Sterli et al. 2015; Joyce et al. 2016; Joyce 2017; Evers & Benson 2019). The species of *Basilemys* were added as sister taxa onto the stem of *Trionychia*, following the generally accepted position of these turtles (e.g., Brinkman 1998; Joyce et al. 2016). As phylogenetic comparative methods use one specimen per species, we selected the well-preserved AMNH FARB 5448 to represent *Basilemys variolosa*. For extant species, we used species means as in Lichtig and Lucas (2017). We performed 100 pFDA replicates based on the posterior sample of 100 trees from Thomson *et al*. (2021). This procedure allowed us to obtain a distribution of posterior probabilities (PPs) of predicted habitats for the fossils. In all cases, all categories were assigned equal prior probability, as this decreases posterior misidentification rates (Motani & Schmitz 2011). The robustness of the predictions was analyzed based on a two-factor rationale, inspecting (i) the success rates of predicting known ecological categories for extant turtles, whereby we label success of >75% as good and >90% as very good; and the (ii) the median PPs of each predicted habitat, whereby a PP > 75% is considered high and PP>90% very high. This is conservative, as any PP > 50% indicates the presence of an estimated trait. The success rates are assessed for each ecological trait that is estimated, and overall success rates are also computed. In addition, for the prediction of fossils, we counted the number of times each taxon was predicted as having a specific habitat (e.g., Fabbri et al. 2022a), whereby high numbers indicate consistent predictions regardless of the posterior probability. All supplementary data and R scripts are provided in GitHub (https://github.com/G-Hermanson/Reply_shell_measurements_LL2017).

## Results

### PCA results: Overlap in ecologies and changes in morphospace occupation

Biplots of shell doming vs. relative plastron width using the corrected dataset of Lichtig and Lucas (2017) show that terrestrial turtles, on average, have high domed shells and relatively broad plastra, whereas aquatic turtles, in general, show the opposite relationship. This is true regardless of whether shell doming is measured sagittally (i.e., as carapace length–height ratio; Fig. 1A, C) as was done by Lichtig and Lucas (2017), or by the more commonly used coronal doming (i.e., carapace width–height ratio; Fig. 1B, D). Overall, extant and fossil species have similar distributions in all plots, indicating that the choice of how to measure doming has little impact at least on the graphical distribution of data points. The area of ecological overlap has nearly the same extent along the doming axes in the plots that use species means (Fig. 1C–D) and in the plots using specimen-level data (Fig. 1A–B), although the species mean data causes a lesser density of high-domed aquatic turtles. This pattern of reduced density is caused by only a few species, such as *Cuora amboinensis*, for which the measured specimens have a relatively large range of doming values.

**Figure 1.**
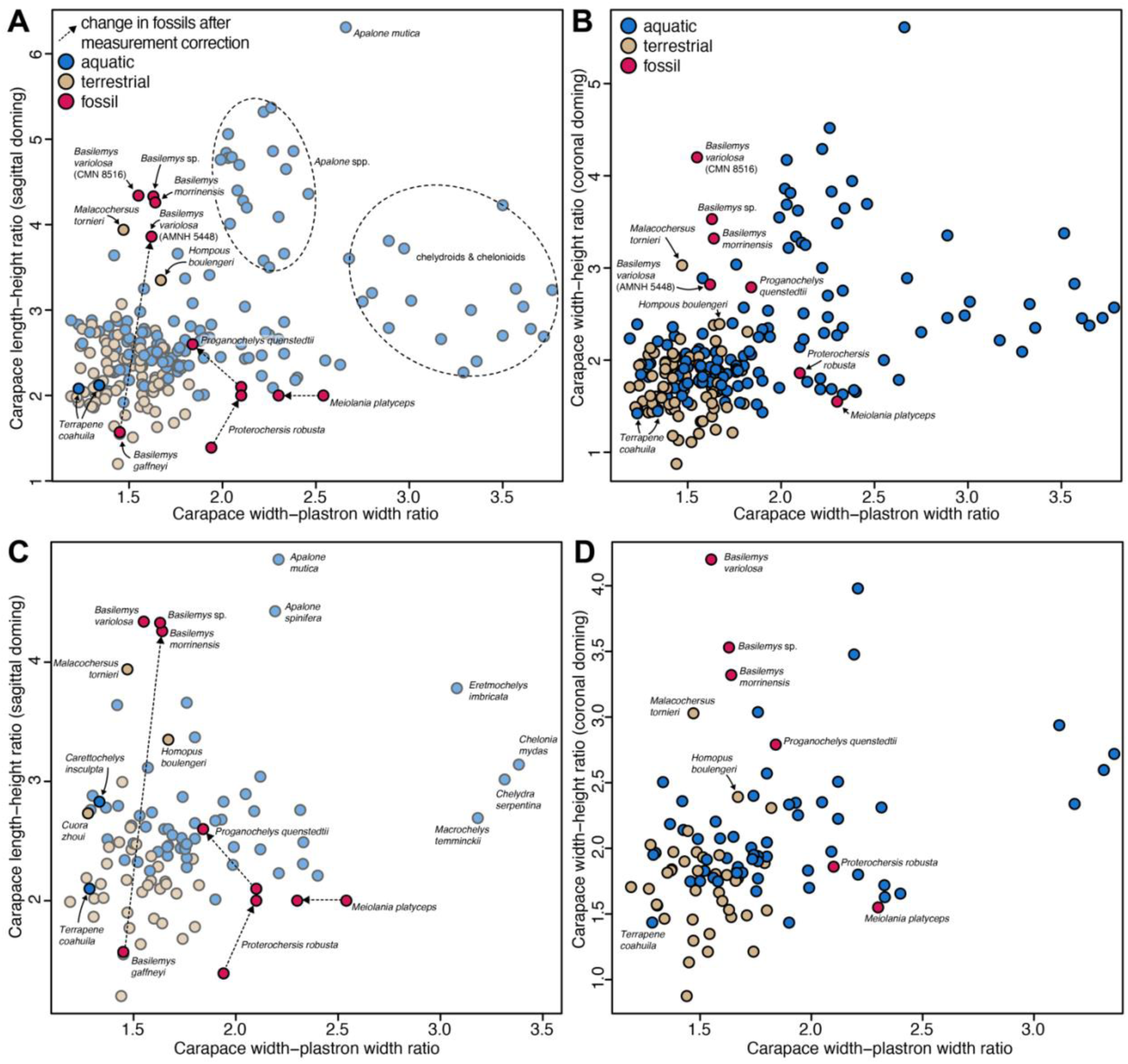
Plots of shell doming against relative plastral width. **A**, using sagittal doming, as in Lichtig and Lucas (2017), on the specimen level. **B**, using coronal doming, on the specimen level. **C**, using sagittal doming, as in Lichtig and Lucas (2017), but using species means for species with multiple individuals. **D**, using coronal doming, using species means. Note that most datapoints in A and C are shown as transparent, to highlight the positions of fossils and previously omitted extant taxa as well as some species discussed in the main text. Fossil datapoint pairs represent original (i.e., uncorrected) values provide by Lichtig and Lucas (2017) and corrected versions of the same species or closely related species in the case of *Basilemys* (see methods), for which the arrow shows the shortest possible change in morphospace. A version of this plot with numbers that correspond to species, and a key can be found in Supplementary Figure 1.

The ecological distribution of data points in all of our corrected plots is consistent with the general pattern reported by Lichtig and Lucas (2017). However, our plots show a graphically much larger area of overlap between ecological groups of turtles (Fig. 1A, C). This is mostly attributable to the fact that the revised dataset includes all explicitly or non-explicitly removed outliers, including the terrestrial *Malacochersus tornieri* and *Homopus boulengeri*, which plot among aquatic turtles, or the aquatic *Terrapene coahuila*, which plots among terrestrial turtles. Nevertheless, the large area of overlap between aquatic and terrestrial turtles remains even if these species are disregarded. For example, the highly aquatic *Carettochelys insculpta* has a shell doming that is nearly identical to that of the terrestrial geoemydid *Cuora zhoui* in the sagittal doming plot (Fig. 1C). Despite the large overlap of ecologies, some regions of the morphospace are exclusively occupied by aquatic turtles. For example, species of the trionychid softshell genus *Apalone* strongly expand the morphospace along high values of the doming axis (i.e., extremely low shell morphologies). The morphospace expansion of aquatic turtles toward low relative plastron widths is caused by chelydrids (i.e., *Macrochelys temminckii* and *Chelydra serpentina*) and chelonioids (i.e., *Chelonia mydas* and *Eretmochelys imbricata*) (Fig. 1).

Correction of measurements for the fossils we retained from the original study (Lichtig & Lucas 2017) also changes their positions among the data population for the sagittal doming plots (Fig. 1A, C). *Proterochersis robusta* moves from a position close to predominantly terrestrial species into an area of the plot where exclusively aquatic taxa are found, caused by its low relative plastron width. *Proganochelys quenstedtii* moves along the doming axis into a region of the plot with intermediate levels of doming. Although predominantly surrounded by aquatic turtles, *Proganochelys quenstedtii* plots at the edge of the area of greatest ecological overlap. *Meiolania platyceps* plots in a position along the relative plastral width axis that is otherwise only populated by aquatic turtles, consistent with Lichtig and Lucas (2017). Our datapoints for *Basilemys morrinensis*, *Basilemys variolosa* and the indeterminate specimen of *Basilemys* sp. plot in complete opposite ends of the doming axis of the plot than the *Basilemys gaffneyi* datapoint of Lichtig and Lucas (2017), highlighting the errors associated with the *Basilemys gaffneyi* measurement provided in Lichtig and Lucas (2017). Variation among the four novel *Basilemys* specimens included could represent individual, taxonomic or taphonomic differences in relative doming. Taphonomic differences are also supported by the observation that the four *Basilemys* specimens show larger variation in coronal doming than sagittal doming, suggesting stronger carapace width differences between specimens than carapace length differences, a pattern that can likely be attributed to taphonomy. Nevertheless, given the overall good preservation state of these fossils, our datapoints indicate the low doming shape of the carapace that has also been reported for the genus in general (e.g., Langston 1956; Sullivan et al. 2013; Mallon & Brinkman 2018). Although this region of the plot is predominantly populated by aquatic turtles, it also includes terrestrial tortoises with relatively low domed shells (i.e., *Malacochersus tornieri* and *Homopus boulengeri*), which had been omitted by Lichtig and Lucas (2017).

### Analysis of linear shell measurements

The data point distributions in our biplots (Fig. 1) should, in our view, not be seen as hypothesis tests of the ecologies of these turtles, which can only be done if the discriminating strength of the relationships depicted are tested with phylogenetic comparative methods. Our respective phylogenetic Flexible Discriminant Analysis indicates that the two ecological categories (i.e., aquatic and terrestrial) cannot be differentiated based on the relationships of sagittal or coronal doming and relative plastral width. This even holds true when the highest-domed aquatic turtles of the dataset (i.e., *Terrapene coahuila*) and the flattest-shelled terrestrial turtles (i.e., *Malacochersus tornieri* and *Homopus boulengeri*) are excluded from the dataset. Although success rates for predicting aquatic ecology are high (for sagittal doming consistently 95.8% across 100 trees using the full dataset and a median value of 95.7% using the reduced dataset; for coronal doming consistently 95.8% across 100 trees using the full dataset and 95.7% using the reduced), this is contrasted by extremely low success rates for predicting terrestrial ecology for sagittal doming (median of 3.6% across 100 trees using the full dataset and 7.4% using the reduced dataset) as well as for coronal doming (consistently 0 across 100 trees using the full dataset and a median of 3.7% using the reduced). This amounts to an overall median success rate for predicting the correct ecology of 64.5% for sagittal doming in the full dataset (63.9% in the reduced dataset) and 63.1% for coronal doming in the full dataset (64.8% in the reduced dataset), but also means that every turtle is predicted to be aquatic, which is not very helpful. This is confirmed by all five fossil species being predicted to be aquatic, with high posterior probabilities for both sagittal doming (*Proganochelys quenstedtii* PP_median_= 99.1%; *Proterochersis robusta* PP_median_ = 98.9%; *Meiolania platyceps* PP_median_ = 78.5%; *Basilemys morrinensis* PP_median_ = 85.1%; *Basilemys variolosa* PP_median_= 85.5% across 100 trees) and coronal doming (*Proganochelys quenstedtii* PP_median_= 99.6%; *Proterochersis robusta* PP_median_ = 99.4%; *Meiolania platyceps* PP_median_ = 80.0%; *Basilemys morrinensis* PP_median_ = 81.4%; *Basilemys variolosa* PP_median_= 85.8% across 100 trees) when using the full dataset. Using the reduced dataset, results are very similar (sagittal doming: *Proganochelys quenstedtii* PP_median_= 99.0%; *Proterochersis robusta* PP_median_ = 98.3%; *Meiolania platyceps* PP_median_ = 76.4%; *Basilemys morrinensis* PP_median_ = 88.7%; *Basilemys variolosa* PP_median_= 89.3% across 100 trees; coronal doming: *Proganochelys quenstedtii* PP_median_= 99.7%; *Proterochersis robusta* PP_median_ = 99.3%; *Meiolania platyceps* PP_median_ = 78.2%; *Basilemys morrinensis* PP_median_ = 84.8%; *Basilemys variolosa* PP_median_= 90.4% across 100 trees). Given that the data cannot discriminate between extant aquatic and terrestrial species, the high posterior probabilities for fossil predictions are rendered meaningless, and it must be concluded that the shell measurements used in this study cannot predict ecology in turtles, whereby it does not matter if shell doming is quantified along the long axis or transverse width of the turtle carapace.

## Discussion

When examining the Lichtig and Lucas (2017) dataset, we found numerous reproducibility issues. Correcting fossil species measurements, correcting some ecological classifications (e.g., *Malachersus tornieri* was corrected to be terrestrial), including all datapoints (aquatic turtles with high-domed carapaces such as *Terrapene coahuila* were reinserted), and using adequate phylogenetic discriminant analysis returns strongly different results that no longer support the central claims of the Lichtig and Lucas (2017) study. Based on our results, the simple carapace measurements proposed by Lichtig and Lucas (2017) are insufficient to characterize or infer ecology of turtles. This is not only because several extant aquatic and terrestrial turtle species exist that show doming that is atypical for the majority of their ecological guild, as our analyses excluding these species demonstrate. More importantly, this is because there is a large region of morphological overlap along the gradient of doming. Using finer habitat ecologies does not lead to a gradational pattern of doming according to habitat ecology: Our Figure 2 shows species mean data for the recorded shell ratios, whereby species are colour-coded according to their intensity of hand-webbing, which is a morphological proxy for their swimming capabilities (e.g., Foth et al. 2019; Dziomber et al. 2020). Although we do not provide formal statistical analyses for this, the webbing-groups show large overlap especially along the doming axis. Our results do not mean that there is no habitat information in turtle shell shape, but that the simple measurements proposed by Lichtig and Lucas (2017) do not work as sufficient discriminators between the two principal habitat ecologies of turtles.

**Figure 2.**
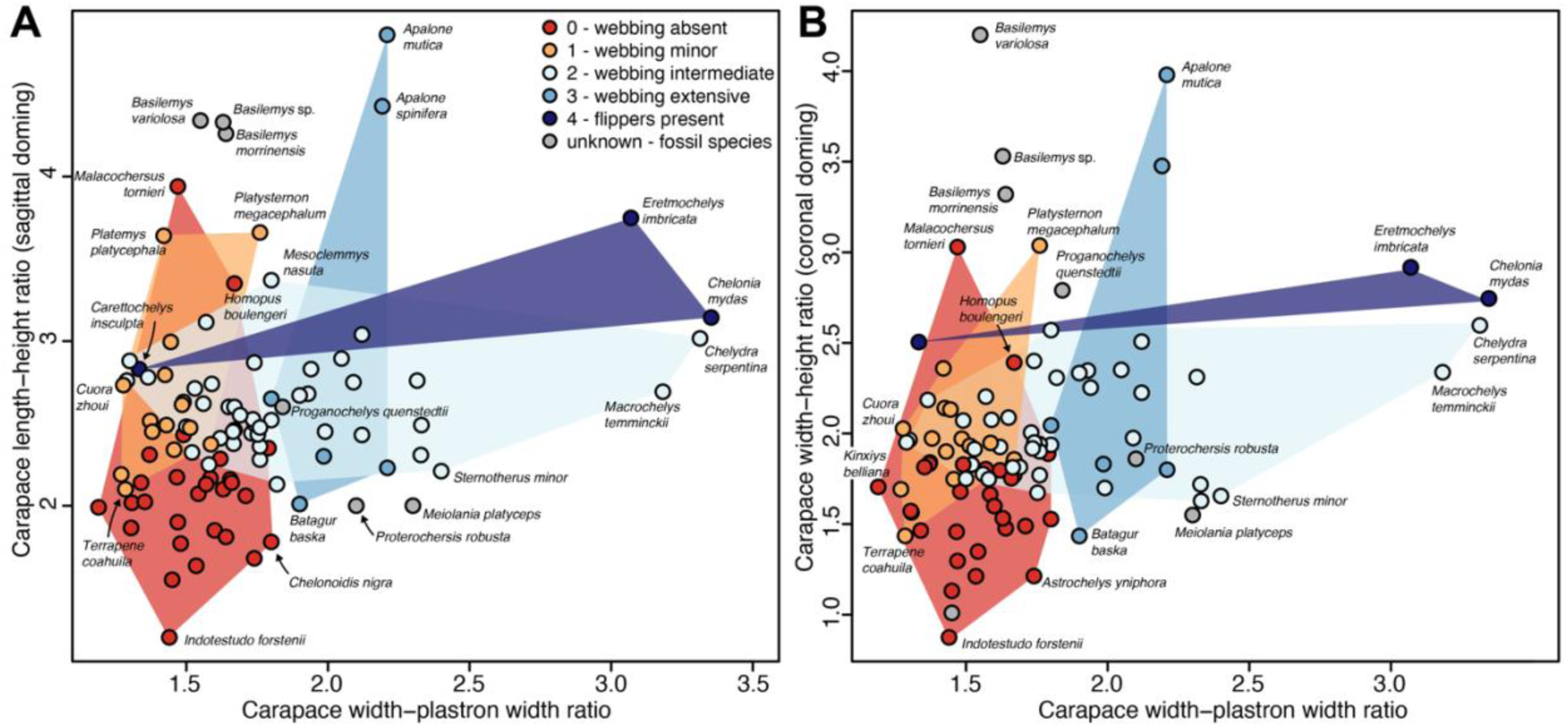
Plots of shell doming against relative plastral width using species means and hand-webbing as an alternative ecological proxy. **A**, using sagittal doming, as in Lichtig and Lucas (2017), using species means. **B**, using coronal doming, on the species means. Fossil datapoint pairs represent corrected or new measurements provided in this study. A version of this plot with numbers that correspond to species, and a key can be found in Supplementary Figure 2.

Although *Proganochelys quenstedtii* plots in an area of the morphospace that is predominantly occupied by aquatic turtles, this should not be misinterpreted as evidence for an aquatic ecology of this fossil species, as was done by Lichtig and Lucas (2017). Contrary to Lichtig and Lucas (2017), *Proterochersis robusta* has no shell dimensions that are uniquely found in terrestrial turtles. The shell shape of *Basilemys* spp. is consistent with aquatic turtles or flat terrestrial turtles, contrary to Lichtig and Lucas (2017) who portray the species as one of the highest domed turtles. *Meiolania platyceps* has shell doming values that are within the range of greatest overlap between aquatic and terrestrial species, but it has a narrow plastron that today is largely observed among aquatic chelydrids and chelonioids, leading Lichtig and Lucas (2017) to the unusual interpretation that *Meiolania platyceps* was an aquatic turtle (contra, e.g., Sterli 2015; Brown & Moll 2019).

An interesting observation from our data is that all stem turtles included in the dataset (i.e., *Proganochelys quenstedtii*, *Proterochersis robusta*, *Meiolania platyceps*) have relatively narrow posterior plastral lobes, although they are most commonly interpreted to be terrestrial (e.g., Joyce 2017). This seems to be the case also in early stem turtles not included in the dataset, such as *Kayentachelys aprix* (a mesochelydian stem turtle: e.g., Sterli & Joyce 2007), *Naomichelys speciosa* (a helochelydrid stem turtle: e.g., Joyce et al. 2014; see phylogeny of Rollot et al. 2022), and *Mongolochelys efremovi* (a sichuanchelyid stem turtle: e.g., Joyce et al. 2016). These species have carapace width to plastron width ratios of 1.86, 2.11, 1.94, respectively, such that they would fall further to the right on the relative plastron width axis than the majority of datapoints in our Figure 1. Thus, it is possible that narrow plastra are plesiomorphic for stem turtles. The evolution of narrow plastra should be researched more, especially because the phylogenetic uncertainty surrounding the stem lineages of americhelydians (i.e., chelonioids+chelydroids) and cryptodires more widely (e.g., Zhou & Rabi 2015) means that it is currently not clear if the narrow plastra of some aquatic cryptodires are symplesiomorphic retentions or independent acquisitions of this trait. For example, an independent origin of narrow plastra could have taken place under different selection pressures, whereas the retaining of the feature could mean that narrow plastra in aquatic cryptodires are either not functionally selected against, or an exaptation (Gould & Vrba 1982). Although this is speculative, the initial narrowness of turtle plastra may have been lost during later turtle evolution due to predation pressure of small mammals, which has also been hypothesized to cause the reduction of skull emarginations among post K/Pg baenid turtles (Lyson & Joyce 2009). As already described by Lichtig and Lucas (2017), the narrower gap between plastron and carapace in extant terrestrial turtles shields the soft tissue parts of these turtles, and this constraint may be relaxed in the bottom-walking chelydrids and marine chelonioids (Hermanson et al. 2024).

We strongly advocate researchers not to use plots that are intended to show general distribution patterns of data as a substitute to statistical hypothesis tests, or as an equally informative tool for data interpretation. Plots (biplots as here, or PCA morphospaces) can aid in formulating hypotheses, but these should be formally evaluated using appropriate methods that account for the peculiarities of the data, such as phylogenetic autocorrelation in species data (Felsenstein 1985).

## Institutional abbreviations

AM: Australian Museum, Sydney, Australia.
AMNH FARB: American Museum of Natural History, Fossil Amphibians, Reptiles and Birds, New York City, New York, USA.
CMN: Canadian Museum of Nature, Ottawa, Canada.
DMNS: Denver Museum of Nature & Science, Denver,Colorado, USA.
NHMUK: Natural History Museum, London, UK.
SMF: Senckenberg Museum Frankfurt, Frankfurt, Germany.
SMNS: Staatliches Museum für Naturkunde, Stuttgart, Germany.
UF: University of Florida, Florida Museum of Natural History, Gainesville, Florida, USA.
USNM: United States National Museum, Washington, DC, USA.
YPM: Yale Peabody Museum, New Haven, Connecticut, USA.

## Data, script and code Availability Statement

All R code and primary data tables required to reproduce our analyses and plots are deposited at GitHub (https://github.com/G-Hermanson/Reply_shell_measurements_LL2017) and these data are also archived via zenodo (https://doi.org/10.5281/zenodo.14275543).

## Supplementary information

The documents attached to this manuscript as supplementary information are the following: A data folder that contains (i) a read me file, (ii) spreadsheet with all recorded data, (iii) a tree file for phylogenetic comparative analyses, (iv) two R scripts with which all analyses can be done and with which raw figures can be generated, and (v) a supplementary document that contains versions of figures 1 and 2 with data point numbered according to a species key. These supplementary figures are also provided as separate files (i.e., not embedded in a PDF). These data are available via GitHub (https://github.com/G-Hermanson/Reply_shell_measurements_LL2017) and zenodo (https://doi.org/10.5281/zenodo.14275543).

## Acknowledgements

We thank Roger Benson and Carolyn Merrill for surface scanning and providing access to AMNH 5448, and Ilya Dziomber for discussions. We thank Jordan Mallon and the CMN for uploading CMN FV 57059 to MorphoSource, and Evan Tamez-Galvan and the DMNS for uploading DMNS EPV 103391-4891 to MorphoSource. We thank Heather Smith and Donald Brinkman for reviewing this manuscript, and Jordan Mallon for handling it as an editor at PCI Paleo.

