## Supplementary File 1 for "Simple shell measurements do not consistently predict habitat in turtles: a reply to Lichtig and Lucas (2017)"

This supplementary file contains the species key and by-species labelled version of our Figures 1 and 2 of the main text. Note that PDFs (i.e., searchable files) of Supplementary Figures 1 and 2 are also available in in GitHub (https://github.com/G-Hermanson/Reply_shell_measurements_LL2017).

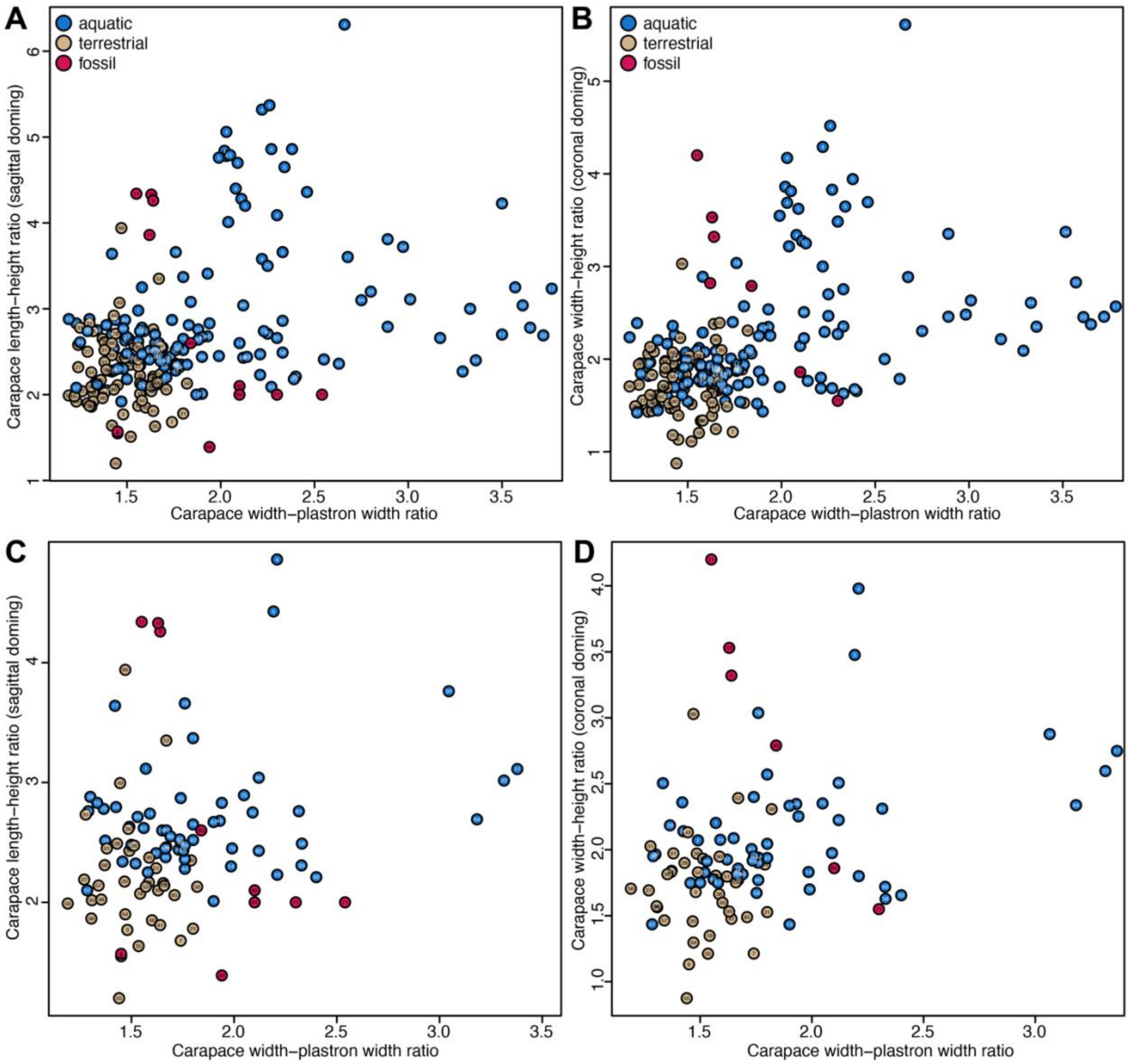


**Supplementary Figure 1.** Plots of shell doming against relative plastral width. **A**, using sagittal doming, as in Lichtig and Lucas (2017), on the specimen level. **B**, using coronal doming, on the specimen level. **C**, using sagittal doming, as in Lichtig and Lucas (2017), but using species means for species with multiple individuals. **D**, using coronal doming, using species means. Species key: 1- *Acanthochelys spixii*; 2- *Actinemys marmorata*; 3- *Agrionemys horsfieldii*; 4- *Apalone mutica*; 5- *Apalone spinifera*; 6- *Astrochelys radiata*; 7- *Astrochelys yniphora*; 8- *Basilemys gaffneyi*; 9- *Basilemys morrinensis*; 10- *Basilemys sp.; 11- Basilemys variolosa*; 12- *Batagur affinis*; 13- *Batagur baska*; 14- *Batagur borneoensis*; 15- *Carettochelys insculpta*; 16- *Chelodina longicollis*; 17- *Chelonia mydas*; 18- *Chelonoidis carbonaria*; 19- *Chelonoidis chilensis*; 20- *Chelonoidis denticulata*; 21- *Chelonoidis nigra*; 22- *Chelydra serpentina*; 23- *Chrysemys picta*; 24- *Clemmys guttata*; 25- *Cuora amboinensis*; 26- *Cuora aurocapitata*; 27- *Cuora galbinifrons*; 28- *Cuora mouhotii*; 29- *Cuora pani*; 30- *Cuora trifasciata*; 31- *Cuora zhoui*; 32- *Dermatemys mawii*; 33- *Emydoidea blandingii*; 34- *Eretmochelys imbricata*; 35- *Geochelone elegans*; 36- *Glyptemys insculpta*; 37-

*Glyptemys muhlenbergii*; 38- *Gopherus agassizii*; 39- *Gopherus berlandieri*; 40- *Gopherus flavomarginatus*; 41- *Gopherus polyphemus*; 42- *Graptemys barbouri*; 43- *Graptemys ernsti*; 44- *Graptemys flavimaculata*; 45- *Graptemys geographica*; 46- *Graptemys nigrinoda*; 47- *Graptemys pseudogeographica*; 48- *Graptemys versa*; 49- *Heosemys depressa*; 50- *Heosemys spinosa*; 51- *Homopus areolatus*; 52- *Homopus boulengeri*; 53- *Homopus femoralis*; 54- *Homopus signatus*; 55- *Indotestudo forstenii*; 56- *Kinixys belliana*; 57- *Kinixys erosa*; 58- *Kinosternon angustipons*; 59- *Kinosternon baurii*; 60- *Kinosternon flavescens*; 61- *Kinosternon scorpioides cruentatum*; 62- *Kinosternon sonoriense*; 63- *Kinosternon subrubrum*; 64- *Macrochelys temminckii*; 65- *Malaclemys terrapin*; 66- *Malacochersus tornieri*; 67- *Malayemys subtrijuga*; 68- *Manouria emys*; 69- *Mauremys nigricans*; 70- *Mauremys reevesii*; 71- *Meiolania platyceps*; 72- *Mesoclemmys gibba*; 73- *Mesoclemmys nasuta*; 74- *Mesoclemmys tuberculata*; 75- *Pelomedusa subrufa*; 76- *Pelusios castaneus*; 77- *Platemys platycephala*; 78- *Platysternon megacephalum*; 79- *Podocnemis expansa*; 80- *Podocnemis lewyana*; 81- *Podocnemis sextuberculata*; 82- *Podocnemis unifilis*; 83- *Proganochelys quenstedtii*; 84- *Proterochersis robusta*; 85- *Psammobates geometricus*; 86- *Pseudemys texana*; 87- *Pseudemys gorzugi*; 88- *Rhinoclemmys areolata*; 89- *Rhinoclemmys funerea*; 90- *Rhinoclemmys pulcherrima*; 91- *Rhinoclemmys rubida*; 92- *Sternotherus minor*; 93- *Sternotherus odoratus*; 94- *Stigmochelys pardalis*; 95- *Terrapene carolina*; 96- *Terrapene carolina triunguis*; 97- *Terrapene coahuila*; 98- *Terrapene ornata*; 99- *Testudo graeca*; 100- *Testudo hermanni*; 101- *Trachemys scripta*.

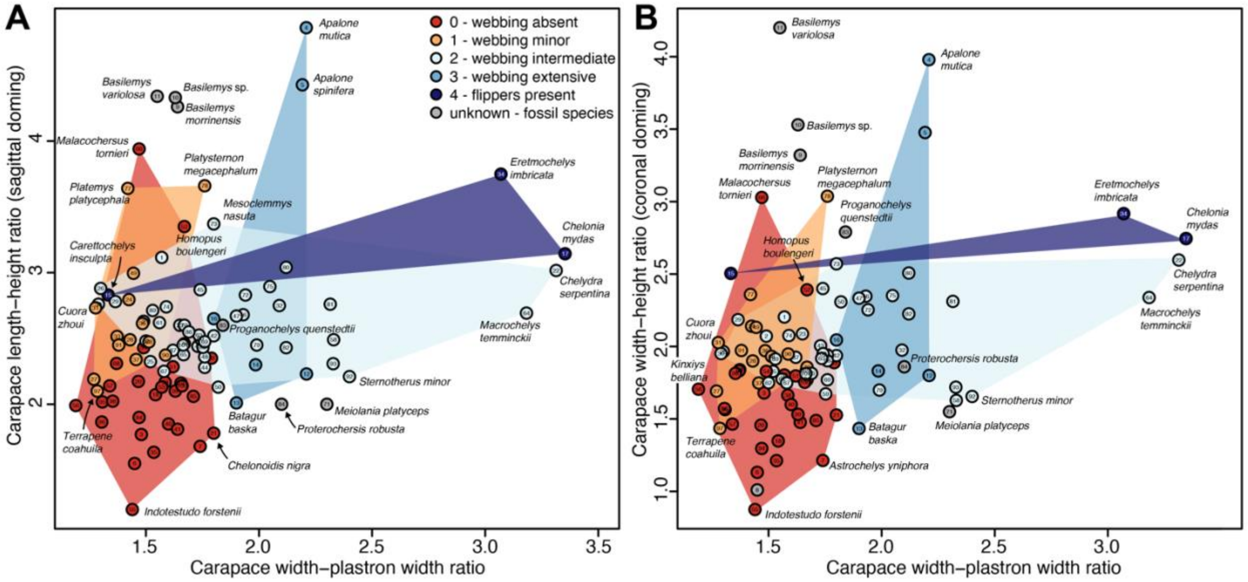


**Supplementary Figure 2.** Plots of shell doming against relative plastral width using species means and hand-webbing as an alternative ecological proxy. **A**, using sagittal doming, as in Lichtig and Lucas (2017), using species means. **B**, using coronal doming, on the species means. Fossil datapoint pairs represent corrected or new measurements provided in this study. Species key as in Supplementary figure 1.
